# miR319-targeted *LsTCP4* and non-target *LsTCP17* act in parallel to promote leaf senescence in lettuce

**DOI:** 10.64898/2026.07.10.737324

**Authors:** Tao Jiang, Sameena Ejaz Tanwir, Sandy Zammar, Kent J. Bradford, Heqiang Huo

## Abstract

Leaf senescence directly affects lettuce quality and postharvest shelf life, but the regulatory roles of miR319-targeted and non-target CIN-TCP transcription factors remain unclear. Here, we examined whether the miR319–TCP module controls lettuce leaf senescence through separable genetic branches. MIR319 overexpression delayed dark-induced senescence, whereas STTM-mediated miR319 suppression accelerated chlorophyll loss, photosynthetic decline, and senescence-marker activation. Disruption of the miR319-targeted gene *LsTCP4* phenocopied MIR319 overexpression, supporting *LsTCP4* as a pro-senescence factor downstream of miR319. We further found that the miR319 non-target CIN gene *LsTCP17* also promoted senescence, as *tcp17* leaves retained more chlorophyll than wild type during dark treatment. Genetic combinations showed that *tcp17* enhanced chlorophyll retention in the OX319 background and partially rescued the accelerated senescence phenotype of S319, indicating that *LsTCP17* acts through a route separable from the miR319-targeted branch. Together, these results reveal a split CIN-TCP architecture in which miR319-targeted *LsTCP4* and non-target *LsTCP17* provide parallel pro-senescence inputs, offering a genetic framework for targeted improvement of lettuce quality.

## Introduction

Leaf senescence is a major determinant of crop productivity, postharvest quality and marketability. In leafy vegetables such as lettuce (*Lactuca sativa*), premature yellowing directly reduces visual quality and shelf life, making the genetic control of senescence highly relevant for both basic biology and crop improvement^1^. The miR319–TCP module is a conserved regulatory module controlling leaf morphogenesis and developmental timing in plants^2,3^. In Arabidopsis, miR319 represses a subset of CIN (*CINCINNATA*)-clade TCP (*TEOSINTE BRANCHED1, CYCLOIDEA*, and *PROLIFERATING CELL FACTOR*) transcription factors, and altered miR319/TCP activity affects leaf shape, jasmonate biosynthesis, maturation and senescence^2–4^. However, the CIN clade also contains non-target members, and evidence linking the miR319–TCP module to senescence has focused predominantly on the miR319-targeted subset. Whether non-target CIN-TCPs independently contribute to leaf senescence, and whether targeted and non-target members act through common or separable regulatory routes, remains unclear, especially in leafy greens where senescence directly determines postharvest value.

In a companion preprint, we identified 33 lettuce TCP genes and classified them into the major PCF (*PROLIFERATING CELL FACTOR*), CIN, and CYC/TB1 (*CYCLOIDEA–TEOSINTE BRANCHED1*) clades^5^. Based on miR319 target prediction and analysis of publicly available degradome data, five CIN-class genes, *LsTCP2, LsTCP3, LsTCP4, LsTCP10* and *LsTCP24*, were identified as high-confidence miR319 targets, whereas *LsTCP13* and *LsTCP17* lacked canonical miR319 target signatures^5^. This target/non-target framework allowed us to test whether miR319-targeted and non-target CIN-TCPs regulate leaf senescence through common or separable routes.

## Results

### *MIR319* overexpression and suppression alter CIN-TCP expression in lettuce

To examine the relationship between miR319 abundance and CIN-TCP transcript accumulation, we generated *MIR319* overexpression plants (OX319) and short tandem target mimic (STTM)-based miR319 suppression lines (S319). Both pri-*MIR319* transcript and mature miR319 abundance were strongly elevated in OX319 relative to wild type (WT) and S319 (Figure 1a). qRT-PCR analysis of selected CIN-TCP genes showed reduced transcript accumulation for several genes in OX319, with *LsTCP4* showing a pronounced reduction (Figure 1b). We therefore selected *LsTCP4* for functional analysis using CRISPR/Cas9 editing. OX319 plants displayed relatively compact architecture with curled or serrated leaves, whereas S319 plants showed a contrasting developmental phenotype. The *tcp4* mutant showed a similar compact growth habit and altered leaf morphology (Figure 1c,d). These results suggest that *LsTCP4* contributes to miR319-associated regulation of lettuce leaf morphology.

**Figure 1.**
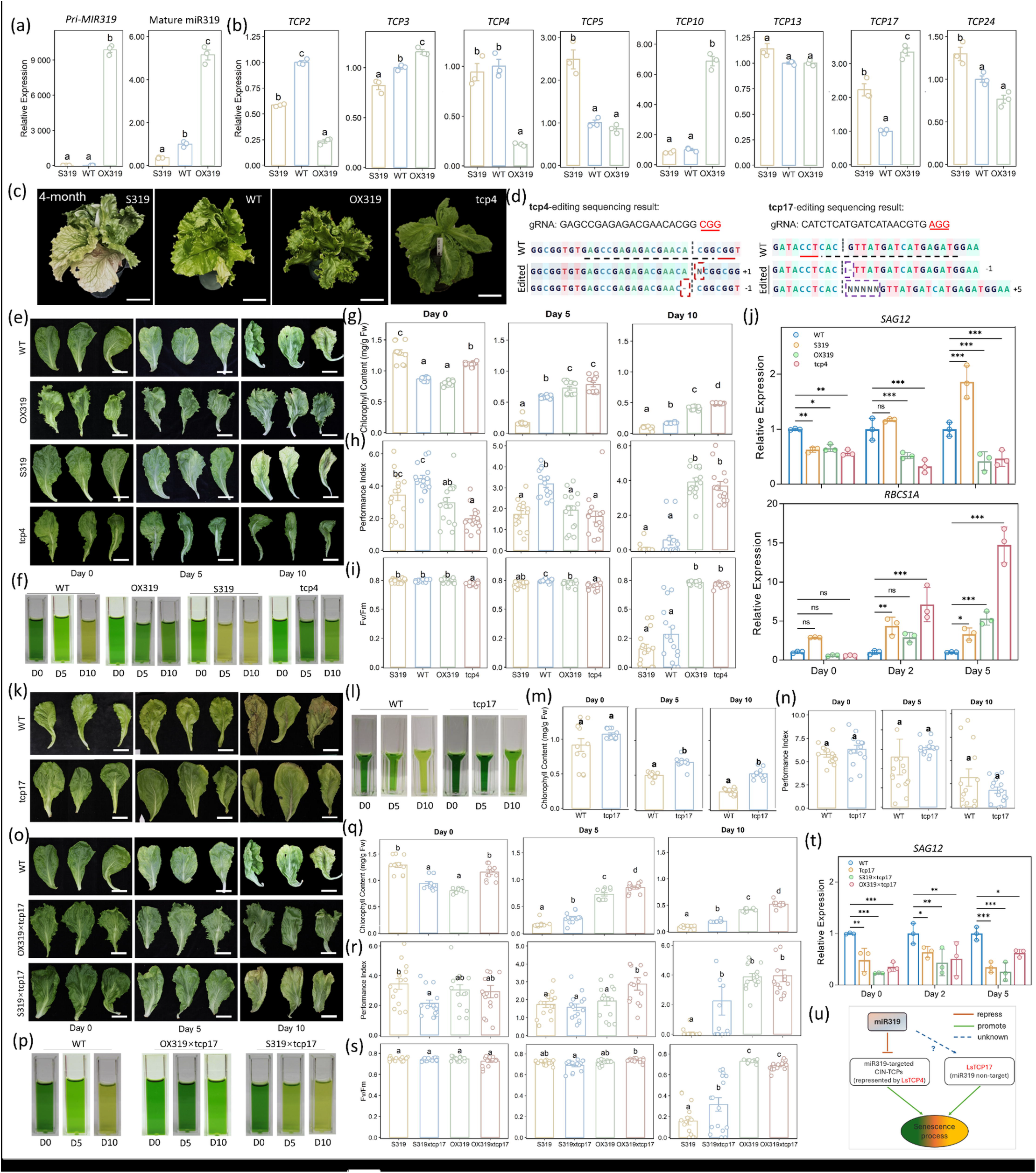
Roles of miR319, *LsTCP4*, and *LsTCP17* in dark-induced leaf senescence in lettuce. **a**, qRT-PCR analysis of pri-*MIR319* transcript and mature miR319 abundance in leaf tissue from WT, *MIR319* overexpression (OX319), and STTM-miR319 suppression (S319) plants. **b**, qRT-PCR analysis of selected CIN-TCP genes in the same leaf samples. **c**, Representative 4-month-old S319, WT, OX319, and *tcp4* plants grown in controlled-environment growth rooms. **d**, CRISPR/Cas9-induced mutations in *LsTCP4* and *LsTCP17*. Target sequences, protospacer adjacent motif (PAM) regions, and edited sites are indicated. **e**, Detached leaves of WT, OX319, S319, and *tcp4* plants during dark-induced senescence (DIS) at days 0, 5, and 10. **f**, Representative chlorophyll extracts from the genotypes shown in e. **g**–**i**, Total chlorophyll content (**g**), photosynthetic performance index (PIABS; **h**), and Fv/Fm (**i**) during DIS. **j**, Relative expression of *SAG12* and *RBCS1A* in S319, WT, OX319, and *tcp4* leaves at days 0, 2, and 5 of DIS. **k**, Detached leaves of WT and *tcp17* plants during DIS at days 0, 5, and 10. **l**, Representative chlorophyll extracts from WT and *tcp17* leaves. **m, n**, Total chlorophyll content (**m**) and PIABS (**n**) in WT and *tcp17* leaves during DIS. **o**, Detached leaves of WT, OX319 × *tcp17*, and S319 × *tcp17* plants during DIS at days 0, 5, and 10; corresponding OX319 and S319 leaves are shown in **e. p**, Representative chlorophyll extracts from the genotypes shown in **o. q**–**s**, Total chlorophyll content (**q**), PIABS (**r**), and Fv/Fm (**s**) in S319, S319 × *tcp17*, OX319, and OX319 × *tcp17* leaves during DIS. **t**, Relative *SAG12* expression in the genotypes shown in **q–s. u**, Proposed working model of the relationships among miR319-targeted CIN-TCPs, represented by *LsTCP4*, and non-target *LsTCP17* during leaf senescence. qRT-PCR data in **a, b, j**, and **t** are means ± SEM of three independent biological replicates. Physiological data in **g–i, m, n**, and **q–s** are means ± SEM of six biologically independent detached leaves per genotype at each time point. Different letters indicate significant differences among genotypes within each time point in panels **g–i, m, n**, and **q–s**. In panels **j** and **t**, horizontal brackets indicate pairwise comparisons; ns, not significant; \**P* < 0.05; \*\**P* < 0.01; \*\*\**P* < 0.001. Scale bars, 5 cm in **c** and 1 cm in **e, k**, and **o**.

### *MIR319* overexpression and *LsTCP4* disruption delay dark-induced leaf senescence

Differences in leaf yellowing observed during routine growth prompted us to test whether this developmental module also regulates senescence, a trait directly tied to lettuce postharvest value. Detached leaves of wild type, OX319, S319, and *tcp4* plants were subjected to dark-induced senescence (DIS) and evaluated at days 0, 5, and 10 (Figure 1e). S319 leaves yellowed earlier and lost chlorophyll faster than wild type, whereas OX319 leaves showed delayed yellowing, and *tcp4* leaves phenocopied OX319, remaining greener during prolonged darkness. The genotype-dependent differences became more pronounced over time, indicating distinct senescence trajectories during dark treatment.

Physiological measurements were performed on detached leaves collected at each DIS time point and further supported this interpretation. Representative chlorophyll extracts showed faster pigment loss in S319 and stronger pigment retention in OX319 and *tcp4* (Figure 1f). Quantification confirmed this trend: at day 10, OX319 retained 141.3% more chlorophyll than wild type, whereas S319 showed 42.6% lower chlorophyll content than wild type (Figure 1g). Chlorophyll fluorescence measurements provided an independent functional readout. Photosynthetic performance index (PIABS) and Fv/Fm, which reflect overall photosynthetic performance and maximum PSII efficiency, declined fastest in S319 but were better maintained in OX319 and *tcp4* during prolonged darkness (Figure 1h, i). At day 10, OX319 maintained 1.6-fold higher Fv/Fm and 6.0-fold higher PIABS than wild type, whereas S319 showed 44.3% lower Fv/Fm and 84.1% lower PIABS. Thus, miR319 overexpression and *LsTCP4* disruption delayed both pigment loss and photosynthetic decline. The marker gene expression results were consistent with this conclusion: expression of the senescence-associated marker *SAG12* was significantly reduced in OX319 and *tcp4*, whereas the photosynthesis-associated marker *RBCS1A* was better maintained (Figure 1j). These results are consistent with *LsTCP4* acting downstream of miR319 as a pro-senescence factor.

### The miR319 non-target gene *LsTCP17* also promotes leaf senescence

The CIN clade also includes members lacking canonical miR319 target signatures, raising the possibility that miR319 non-targeted CIN-TCPs may contribute additional senescence control. *LsTCP17* was of particular interest because it belongs to the CIN clade yet showed no evidence of direct miR319-guided cleavage^5^. CRISPR/Cas9-mediated disruption of *LsTCP17* was confirmed by sequencing (Figure 1d). During DIS, *tcp17* leaves yellowed more slowly than wild type and stayed green longer (Figure 1k). Representative chlorophyll extracts also showed stronger pigment retention in *tcp17* during dark treatment (Figure 1l). Quantitative measurements confirmed delayed senescence in *tcp17*: at day 10, *tcp17* leaves retained twice as much chlorophyll as wild type, although photosynthetic performance parameters were not significantly different from wild type under this assay condition (Figure 1m, n). Thus, *LsTCP17* also promotes senescence, but through a mechanism not explained by direct miR319 cleavage. This finding extends the miR319–TCP framework beyond direct miRNA-target regulation and points to a broader CIN-TCP senescence network.

### *tcp17* genetically modifies OX319 and S319 senescence phenotypes

To determine whether *LsTCP17* acts in the same pathway as miR319-targeted TCPs or in a parallel branch, we introduced *tcp17* into both OX319 and S319 backgrounds. These combinations provided a genetic test of pathway relationships. If *LsTCP17* functioned solely downstream of the miR319-targeted branch, adding *tcp17* to OX319 would be expected to produce little additional delay in senescence. Instead, OX319 × *tcp17* leaves showed stronger greenness retention than OX319 alone during DIS (Figure 1o, p). At day 10, OX319 × *tcp17* retained 25.3% more chlorophyll than OX319 and 200.2% more chlorophyll than wild type (Figure 1g, q). Conversely, introducing *tcp17* into the S319 background partially rescued the accelerated senescence phenotype caused by miR319 reduction, approximately doubling chlorophyll retention (100.4% higher) relative to S319. Chlorophyll fluorescence analysis showed a consistent pattern: OX319 × *tcp17* maintained high PIABS and Fv/Fm, whereas S319 × *tcp17* partially restored PIABS and Fv/Fm relative to S319 (Figure 1r, s). Senescence-associated marker *SAG12* expression further supported this genetic interaction, with *tcp17* reducing the pro-senescence transcriptional output in the S319 background and reinforcing the delayed-senescence state in the OX319 background (Figure 1t).

## Discussion

These genetic interactions support a model in which miR319-targeted *LsTCP4* and non-target *LsTCP17* provide genetically separable inputs to leaf senescence. Rather than acting as interchangeable components of one linear pathway, the miR319–*LsTCP4* branch and non-target *LsTCP17* represent genetically separable inputs that both feed into the senescence process. In this model, miR319 delays senescence by repressing targeted CIN-TCPs such as *LsTCP4*, while non-target *LsTCP17* promotes senescence through a route that is independent of direct miR319 targeting (Figure 1u). Thus, miR319-targeted *LsTCP4* and non-target *LsTCP17* define separable upstream inputs that converge on shared senescence outputs, including chlorophyll degradation, photosynthetic decline and activation of senescence-associated transcriptional programs.

Several limitations should be considered when interpreting these findings. Our assays were conducted primarily under dark-induced senescence conditions using detached lettuce leaves. Although dark-induced senescence is widely used as an experimental proxy to accelerate and synchronize senescence responses^1^, it does not fully reproduce commercial postharvest storage, where cold temperature, light history, humidity, packaging, and microbial pressure may also influence shelf-life performance. This study focused on *LsTCP4* as a representative miR319-targeted CIN-TCP and *LsTCP17* as a non-target CIN-TCP. Other miR319-targeted genes, including *LsTCP2, LsTCP3, LsTCP10*, and *LsTCP24*, may also contribute to senescence regulation, either redundantly or in a genotype- or context-dependent manner. Although the genetic interactions support separable inputs from the miR319–*LsTCP4* branch and *LsTCP17*, the downstream transcriptional targets and hormone-related mechanisms connecting these TCPs to chlorophyll degradation and photosynthetic decline remain to be determined. Future work should evaluate these lines under commercial cold-storage conditions and test additional single and combinatorial CIN-TCP mutants to define the broader regulatory network.

This work reveals a split CIN-TCP architecture controlling lettuce senescence. One branch is miR319-dependent, represented by targeted CIN-TCPs such as *LsTCP4*, whereas the other is miR319 non-targeted and represented by *LsTCP17*. This distinction has practical value for leafy crop improvement. Although miR319 manipulation delays senescence, it also alters leaf morphology, consistent with the broad developmental roles of miR319-regulated TCP factors^2,3^. By contrast, *LsTCP17* may offer a more focused target for testing postharvest shelf-life improvement while potentially avoiding some pleiotropic effects of broad miR319 manipulation. Together, our results identify *LsTCP4* and *LsTCP17* as parallel pro-senescence regulators and provide a genetic framework for targeted improvement of lettuce quality.

## Materials and Methods

### Plant materials and growth conditions

All experiments were performed using lettuce (*Lactuca sativa* L.) in the ‘Salinas’ background. Wild type (WT), MIR319-overexpression plants (OX319), STTM-miR319 suppression lines (S319), CRISPR/Cas9-edited *tcp4* and *tcp17* mutants, and the derived OX319 × *tcp17* and S319 × *tcp17* lines were used in this study.

Seeds were surface-sterilized in 10% (v/v) sodium hypochlorite for 10 min, rinsed seven times with sterile water, and germinated on half-strength Murashige and Skoog medium supplemented with 1% sucrose and 0.8% agar. Seedlings were maintained at 22 °C under a 16-h light/8-h dark photoperiod with a light intensity of approximately 120 μmol photons m^−2^ s^−1^. Seven-day-old seedlings were transferred to a peat-based growth substrate and grown in a controlled-environment chamber at 22 °C/18 °C day/night temperature, 60% relative humidity, a 16-h light photoperiod, and approximately 200 μmol photons m^−2^ s^−1^. Plants used for senescence assays were age-matched and selected for uniform morphology.

### Vector construction and lettuce transformation

For *MIR319* overexpression, the lettuce *MIR319* precursor sequence was amplified and cloned into a binary vector under the control of a constitutive CaMV35S-based promoter to generate OX319 plants. For miR319 suppression, a short tandem target mimic construct targeting miR319 was designed according to the STTM strategy^6^. The STTM-miR319 cassette contained two imperfect miR319-binding sites separated by a 48-nt spacer with a non-cleavable central bulge to sequester endogenous mature miR319 while preventing target cleavage.

For CRISPR/Cas9 editing of *LsTCP4* and *LsTCP17*, guide RNAs targeting coding regions of each gene were designed and cloned into a CRISPR/Cas9 binary vector^7^. All constructs were confirmed by Sanger sequencing and introduced into *Agrobacterium tumefaciens* strain EHA105.

Lettuce transformation was performed using an Agrobacterium-mediated cotyledon explant infection system. Cotyledons from young sterile seedlings were excised and co-cultivated with *Agrobacterium* for 2 d on co-cultivation medium. Explants were then transferred to selection medium supplemented with kanamycin. Regenerated shoots were rooted on selection medium and transferred to soil. Independent transformants were screened by fluorescence and/or PCR-based detection of the transgene. CRISPR-edited lines were genotyped by PCR amplification of the target regions followed by Sanger sequencing. Homozygous or fixed frameshift mutant alleles were carried forward for phenotypic analysis. Target sequences, protospacer adjacent motif (PAM) regions, and edited sites are shown in Figure 1d.

### Generation of OX319 × *tcp17* and S319 × *tcp17* lines

To examine genetic interactions between miR319 dosage and *LsTCP17, tcp17* plants were crossed with OX319 or S319 plants. Progeny were genotyped for the presence of the MIR319-related transgene and the *tcp17* mutation. Lines carrying the corresponding transgene and the *tcp17* mutant allele were selected for dark-induced senescence assays. The OX319 × *tcp17* and S319 × *tcp17* materials were compared with their corresponding OX319 and S319 parental backgrounds in the senescence assays.

### Dark-induced leaf senescence assay

Dark-induced senescence assays were performed using detached fully expanded leaves of comparable developmental age from age-matched plants. Leaves of equivalent developmental rank, typically the third to fourth true leaf from the apex, were excised and placed adaxial-side up under humid conditions in plastic bags or sealed containers to minimize dehydration. Leaves were incubated at 22 °C in complete darkness.

For the WT, OX319, S319, and *tcp4* comparison, detached leaves were evaluated at days 0, 5, and 10 of dark-induced senescence (DIS). For the WT and *tcp17* comparison, detached leaves were evaluated at the same time points. For the crossed-line comparison, detached leaves from WT, OX319 × *tcp17*, and S319 × *tcp17* were evaluated at days 0, 5, and 10; corresponding OX319 and S319 leaves are shown in Figure 1e for comparison. Representative leaf phenotypes were photographed at each time point.

For molecular analysis of senescence-associated marker genes, leaf tissues were collected at days 0, 2, and 5 of DIS as indicated in the figure. For each time point used for qRT-PCR analysis, three independent biological replicates, each collected from an independent plant, were analyzed. Each biological replicate was measured with three technical replicates. Statistical analysis was performed using biological replicate values.

For physiological measurements, including chlorophyll content and chlorophyll fluorescence, six biologically independent detached leaves were analyzed per genotype at each time point, consistent with the Figure 1 legend.

### Chlorophyll extraction and quantification

Total chlorophyll was extracted from approximately 50 mg fresh leaf tissue in 2 mL of 95% (v/v) ethanol. Samples were incubated overnight at 25 °C in darkness with gentle shaking at approximately 40 rpm until pigments were fully extracted. Representative chlorophyll extracts were photographed for visual comparison.

The absorbance of the cleared supernatant was measured at 665 and 649 nm using a spectrophotometer. Chlorophyll a, chlorophyll b, and total chlorophyll concentrations were calculated using the equations of Lichtenthaler and normalized to leaf fresh weight^8, 9^. For the physiological datasets shown in Figure 1g–i, 1m, 1n, and 1q–s, six biologically independent detached leaves were analyzed per genotype at each time point. Each detached leaf was treated as one biological replicate.

#### Chlorophyll fluorescence measurement

Chlorophyll fluorescence parameters were measured from detached leaves at the indicated DIS time points. Leaves were dark-adapted for 30 min before measurement. Fluorescence transients were recorded using a portable chlorophyll fluorometer. Photosynthetic performance index (PIABS) and maximum quantum yield of photosystem II (Fv/Fm) were used to evaluate photosynthetic performance during senescence^9^.

For Figure 1g–i, 1m, 1n, and 1q–s, six biologically independent detached leaves per genotype were measured at each time point. Each detached leaf was treated as one biological replicate.

### RNA extraction, reverse transcription, and qRT-PCR analysis

Total RNA was extracted from approximately 100 mg of leaf tissue using TRIzol reagent or RNAzol RT according to the manufacturer’s instructions, followed by DNase I treatment to remove genomic DNA contamination. RNA concentration and purity were assessed using a spectrophotometer.

For mRNA and pri-MIR319 quantification, first-strand cDNA was synthesized from 1 μg total RNA using reverse transcriptase. For mature miR319 quantification, stem-loop reverse-transcription primers specific to miR319 were used, followed by qPCR with miRNA-specific forward primers and a universal reverse primer. qRT-PCR was performed using SYBR Green-based chemistry.

Relative expression levels were calculated using the 2^−ΔΔCt^ method with internal reference genes. The analyzed transcripts included pri-MIR319, mature miR319, selected CIN-TCP genes, and senescence-associated marker genes, including *SAG12* and *RBCS1A*. qRT-PCR analyses shown in Figure 1a, 1b, 1j, and 1t were performed using three independent biological replicates. Each biological replicate was measured with three technical replicates, and statistical analyses were performed using biological replicate values.

### Statistical analysis

Data are presented as means ± SEM, consistent with the Figure 1 legend. For qRT-PCR analyses in Figure 1a, 1b, 1j, and 1t, three independent biological replicates were analyzed. For physiological measurements in Figure 1g–i, 1m, 1n, and 1q–s, six biologically independent detached leaves were analyzed per genotype at each time point.

For comparisons among multiple genotypes within the same time point, one-way analysis of variance followed by Tukey’s multiple-comparison test was used. Different letters indicate significant differences among genotypes within each time point in Figure 1g–i, 1m, 1n, and 1q–s. For qRT-PCR panels with planned pairwise comparisons, horizontal brackets indicate pairwise comparisons. Statistical significance was defined as ns, not significant; \**P* < 0.05; \*\**P* < 0.01; \*\*\**P* < 0.001.

## Acknowledgments

This work was supported by the USDA-NIFA grant 2019-67013-29236, the USDA HATCH program FLA-MFC-006387, and FFAR grant A21-1857-S001, awarded to H.H.

## Author contributions

T.J. conceived the study, performed the experiments and data analysis, and drafted the manuscript. S.E.T. and S.Z. contributed to plant materials, phenotyping, and molecular analyses. H.H. supervised the study. K.J.B. contributed manuscript input and funding support. All authors reviewed and approved the manuscript.

## Data availability

The data supporting the findings of this study are available from the corresponding author upon reasonable request.

